# Sound–shape correspondences in macaques reveal evolutionary roots of sound symbolism

**DOI:** 10.64898/2026.07.31.741992

**Authors:** Maria Loconsole, Chunyun Xue, Elias Garcia-Pelegrin

## Abstract

Humans reliably associate certain speech-like sounds with visual shapes, most notably in the Bouba–Kiki Effect, where rounded and spiky shapes are matched with the sounds Bouba and Kiki, respectively. Although often linked to language and culture, evidence from preverbal infants and domestic chickens suggests that sound–shape correspondences may reflect an experience-independent perceptual bias. However, studies on great apes failed to detect such bias, leaving open the question on its phylogenetic origin. Using a free-choice task, we showed the Bouba-Kiki effect in nine macaques, thus suggesting that such correspondences represent a conserved feature of vertebrate perception that may have provided a scaffold for the later emergence of symbolic communication systems in our species.

## INTRODUCTION

Humans exhibit a robust tendency to associate particular speech-like sounds with visual shapes, most famously in the Bouba–Kiki effect, where round shapes are preferentially matched with the non-words Bouba or Maluma, and spiky shapes with Kiki or Takete (Köhler, 1970; Ramachandran & Hubbard, 2001). Despite its robustness, the origins of sound–shape correspondences remain debated (Spence, 2011). Some accounts attribute the Bouba–Kiki effect to experience-dependent factors, such as early exposure to multisensory regularities or to orthographic properties of written language (Lockwood & Dingemanse, 2015). Other accounts propose that it reflects an experience-independent perceptual bias, potentially constituting a predisposed mechanism that facilitates early communication and language acquisition (Ozturk et al., 2013).

Separate evidence comes from newly hatched domestic chickens, which spontaneously match Bouba and Kiki sounds to rounded and angular shapes, respectively, despite minimal and experimentally controlled sensory experience (Loconsole et al., 2026). Although birds are evolutionarily distant from primates, these findings provide compelling support for an experience-independent origin of sound–shape correspondences. Results from chicks challenge the notion that the Bouba–Kiki effect is solely a language-related phenomenon, instead suggesting that it reflects a fundamental crossmodal correspondence (Spence, 2011) that may arise from natural properties of objects and sounds (Fort & Schwartz, 2022). Yet, from an evolutionary perspective, the origins of spontaneous sound–shape correspondences remain unclear. Non-human primates, given their close phylogenetic relationship to humans, provide a key test case for determining whether such correspondences are evolutionarily conserved or arose convergently in distant lineages (Emery & Clayton, 2005; Jarvis et al., 2005). Past studies in great apes have failed to report evidence of the Bouba–Kiki effect (Margiotoudi et al., 2019, 2022). However, these studies often relied on extensive training or tasks involving learned auditory–visual pairings, which may tap symbolic or associative processes rather than reveal inherent perceptual biases. Whether non-human primates exhibit spontaneous sound–shape correspondences under conditions that minimize training and symbolic learning remains an open question.

To address this, we tested one pig-tailed macaque (*Macaca nemestrina*) and eight long-tailed macaques (*Macaca fascicularis*) using a task designed to minimize learning and multimodal or symbolic associations, adapting the methodology previously employed in newly hatched chicks. Monkeys first learned to lift the lid of one of two cups to obtain a hidden food reward, with the rewarded cup being signalled by a black shape (with both spiky and round edges) painted on the lid (**Fig. 1A**). Importantly, no auditory stimulation was presented at this stage. At test, both lids displayed a shape (one round, and one spiky), and a repetition of Bouba or Kiki sound was played in the background. Each monkey completed 24 trials divided into short blocks of three, with the position of the round and spiky shapes, as well as Bouba and Kiki playback, counterbalanced between trials. (**Fig. 1B** and **1C**).

**Fig. 1.**
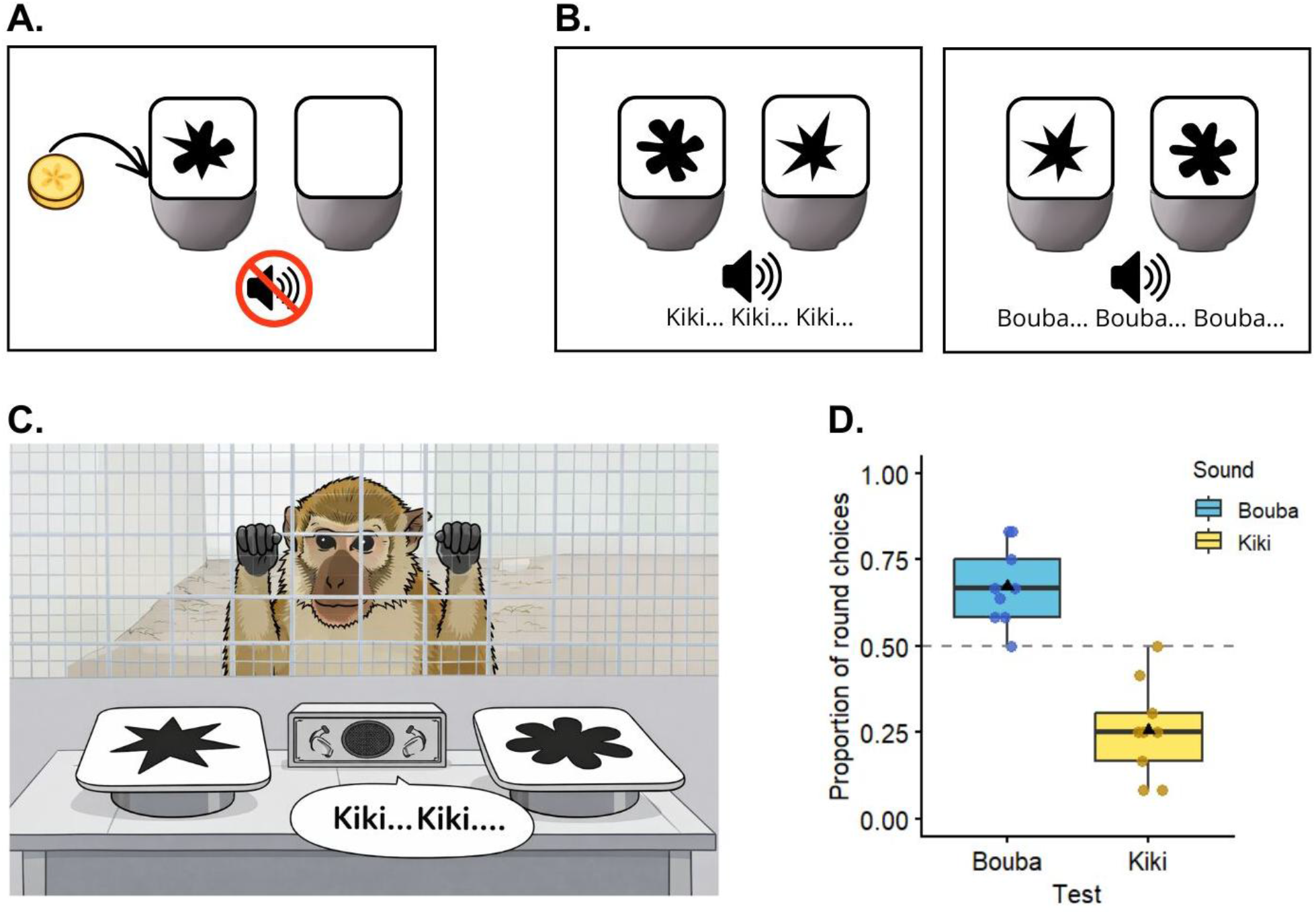
Spontaneous sound–shape correspondences in long-tailed macaques. **A**. Training phase: monkeys learned to lift the lid of one of two cups to obtain a reward. Only the cup with a shape (ambiguous, combining round and spiky edges) contained a reward; no sound was played. **B**. Test phase: both cups displayed distinct shapes (round vs. spiky), and either Bouba or Kiki sounds were played in the background while no reward was provided. **C**. Illustration of a macaque facing the choice task during a Kiki trial. **D**. Proportion of round-shape choices during Bouba and Kiki trials. Monkeys selected the round shape significantly more often during Bouba trials and the spiky shape during Kiki trials. The boxplot’s black bar indicates the median, the black triangle the mean. The dashed gray line indicates chance level; individual data points are shown.

Critically, during training the animals had learned a single rule based on visual shape–reward associations, and no auditory information was available. At test, this rule was no longer applicable, as both options were equally associated with a shape, leaving the animals without an explicitly trained correct response. In this context, we hypothesised that choice behaviour would not reflect learned reinforcement contingencies but could instead be influenced by any pre-existing cross-modal bias. Specifically, if macaques possess a spontaneous sensitivity to sound–shape correspondences, their choices would be biased toward shapes congruent with the concurrently presented sound, similar to humans and chicks.

## RESULTS

The dataset generated during the study is available in **Supplementary Material Data S1**. Analysis using a logistic mixed-effects model revealed a significant effect of sound on shape choice (χ^2^ = 35.14, df = 1, p < .001). Estimated marginal means confirmed that macaques reliably matched sounds to shapes in a manner consistent with the Bouba–Kiki effect (**Fig. 1D**). During Bouba trials, monkeys preferentially chose the round shape (probability = 0.67 ± 0.05 SE), whereas in Kiki trials they selected the spiky shape (probability = 0.74 ± 0.04 SE; both vs. chance, p < 0.001), with monkeys being nearly six times more likely to select the round shape in the Bouba trials (odds ratio = 5.95, 95% CI [∼3–12]).

## DISCUSSION

Our findings demonstrate that macaques spontaneously associate round and spiky shapes with Bouba and Kiki sounds, respectively, under conditions in which the auditory cue is irrelevant and prior learned visual rules no longer provide guidance. This suggests that such crossmodal correspondences can be expressed without explicit training or task-specific learned multimodal associations. A similar effect has previously been observed in chimpanzees for spontaneous pitch-luminance association, indicating that primates can exploit task-irrelevant information when it aligns with crossmodal congruence (Ludwig et al., 2011). Previous evidence of primates relying on crossmodal associations in communicative behaviours further supports the idea that a set of shared associations that links inputs across different sensory modalities can serve an adaptive purpose and ease transmission of information (Ghazanfar et al., 2002; Seyfarth et al., 1980). Thus, rather than being a byproduct of language or culture, these perceptual biases may constitute a fundamental aspect of vertebrate perception, providing a scaffold for the later development of symbolic communication in our species.

Notably, while our findings, as well as previous studies in non-human primates, are based on adult subjects and therefore cannot directly address ontogeny (Spence & Deroy, 2012), evidence from newly hatched chicks (Loconsole et al., 2021, 2026) and preverbal human infants (Mondloch & Maurer, 2004; Ozturk et al., 2013) suggests that at least some crossmodal correspondences emerge very early in development, with limited opportunity for extensive learning. Considering this comparative evidence, a parsimonious interpretation is that these associations may rely, at least in part, on shared predispositions across species. Indeed, it appears less likely that similar crossmodal biases would emerge through entirely different mechanisms in distantly related taxa, such as birds, humans, and non-human primates, with predisposed origins in some species but exclusively experience-dependent learning in others. Rather, the convergence of findings across taxa may point to a common underlying mechanism. Nonetheless, this does not exclude a role for experience. Rather, predispositions may guide organisms toward biologically relevant or frequently occurring regularities in the environment, while experience may subsequently shape, sharpen, or modify these associations over development (Loconsole & Regolin, 2023). Indeed, spontaneous crossmodal associations have been shown to be flexible, as they are subject to extinction in chicks (Loconsole et al., 2021) and tortoises (Loconsole et al., 2023), and they appear to depend on relative rather than absolute stimulus properties in humans (e.g., higher versus lower pitch, rather than absolute pitch values (Brunetti et al., 2018; Spence, 2019)). Together, these findings suggest that experience can modulate crossmodal mappings without necessarily excluding an underlying predisposition.

This framework may also help explain why our monkeys spontaneously expressed the Bouba-Kiki correspondence specifically under conditions of uncertainty, in which previously trained visual rules no longer provided reliable guidance. Conversely, it may indirectly account for inconsistent findings in highly overtrained primates (Margiotoudi et al., 2019, 2022), where extensive training may have overridden or masked spontaneous biases. From an ethological perspective, such predisposed yet flexible mechanisms could provide an adaptive advantage by guiding early interactions with biologically relevant aspects of the environment, while remaining open to modification through experience (Loconsole & Regolin, 2023). Methodological differences between studies may also help contextualise divergent findings in the literature. For instance, in previous studies on great apes, reward delivery during test trials was not contingent on response accuracy but followed a random schedule, which, while methodologically appropriate to avoid reinforcing specific associations, may represent a confounding factor for the subjects that were accustomed to receive reliable feedback on their choices in previous tests. Under such conditions, subjects may adopt a simpler response strategy with reduced sensitivity to the stimuli. Consistent with this interpretation, in the case of Kanzi, the authors reported differences in reaction times between standard training trials and Bouba–Kiki test trials, with slower reaction times in the former (possibly reflecting more deliberative stimulus processing) and faster reaction times in the latter (possibly reflecting reduced engagement and a shallower level of analysis).

Overall, the presence of Bouba–Kiki-like sound–shape matching in humans, naïve chicks, and now macaques raises the possibility that the underlying mechanism is phylogenetically deep, perhaps predating the split between the mammalian and avian lineages. Newly hatched chicks show the same mapping without relevant learning, which argues against a purely language-specific or culturally acquired account, and this macaque results extends that pattern to a non-human primate. Taken together, these findings are consistent with the idea that at least some crossmodal correspondences may rest on ancient perceptual biases inherited from a distant common ancestor, rather than having evolved only within humans. At the same time, the comparative evidence is still too sparse to distinguish firmly between deep homology and convergent evolution. Even so, the joint evidence from chicks and macaques shifts the evolutionary question away from whether the effect is uniquely human, and toward how broadly distributed across vertebrates these perceptual mappings are.

### Limitations and future directions

Because our study was conducted on adult subjects, the present data cannot directly address the ontogenetic origins of the Bouba-Kiki effect. Although the spontaneous nature of the association, together with previous evidence in newly hatched chicks and preverbal infants, is consistent with the existence of predisposed crossmodal heuristics, the relative contribution of innate predispositions and experience-dependent learning remains to be clarified.

Future studies should also investigate more directly the mechanisms through which environmental experience may shape these correspondences across development. Our study does not determine how exposure to environmental regularities may shape, strengthen, attenuate, or modify spontaneous crossmodal mappings over time. Previous evidence suggests that such correspondences are flexible and experience-dependent to some extent, but the degree to which individual sensory histories influence the strength of the effect remains unknown (Spence, 2022). For example, individuals living in richer or more variable natural settings may differ from laboratory-housed animals in the magnitude or stability of the effect. Such comparative approaches may help clarify how spontaneous predispositions and statistical learning interact in the organization of crossmodal perception. In this respect, laboratory paradigms directly manipulating congruent and incongruent crossmodal learning may also help disentangle the respective contributions of spontaneous predispositions and experience-dependent plasticity to the emergence of sound–shape correspondences.

Finally, the present findings are limited to a single non-human primate model and should therefore be extended to additional species and ecological contexts in future research. Comparative studies across taxa, developmental stages, and rearing conditions will be necessary to better clarify the evolutionary origins and functional significance of crossmodal correspondences.

## MATERIALS AND METHODS

### Subjects and housing conditions

A total of nine subjects were tested. Eight were long-tailed macaques (*Macaca fascicularis*; seven females), comprising six adults and two juveniles. The remaining subject was a southern pig-tailed macaque (*Macaca nemestrina*), an adult male. All subjects were part of the Mandai Wildlife Reserve collection. Exact ages are unknown, as individuals had either been confiscated from illegal pet ownership or rescued from roadside environments.

The long-tailed macaques were housed in two separate troops across two enclosures, whereas the pig-tailed macaque was housed individually. Each enclosure measured 7.4 m in length, 4.1 m in width, and 3.2 m in height, and contained enrichment objects as well as tree branches and elevated platforms. During testing, each subject was temporarily separated into a familiar testing enclosure measuring approximately 4.3 m in length, 2.8 m in width, and 3.0 m in height. Subjects were tested either individually or in the presence of a conspecific that had already completed the experiment, provided that the conspecific was a subordinate individual that remained at the back of the enclosure and did not attempt to interact with the subject or interfere with the experimental procedures.

Throughout the testing period, macaques were maintained on their regular diet in accordance with Mandai Wildlife Reserve husbandry guidelines, consisting of leafy and root vegetables provided twice daily. Leafy vegetables were given in the morning to ensure motivation during testing, while root vegetables were provided at the end of the day. Testing rewards consisted of sugary fruits drawn from the subjects’ normal dietary allocation to avoid overfeeding. All subjects had ad libitum access to drinking water.

### Experimental setting

All training and testing sessions were conducted in the subjects’ home enclosures. During both training and testing, subjects stood in front of the experimenter and interacted with the experimental stimuli through a larger opening in the mesh grid of the home cage, which allowed them to extend one arm outside the cage and reach the stimuli. The experimenter sat directly in front of the monkey, with a table placed between them on which the stimuli were presented.

Participation was voluntary: monkeys could choose to take part in the task by approaching the front of the enclosure and positioning themselves near the opening in the mesh grid. Subjects were rewarded with a piece of food when approaching the front of the enclosure. At any point, subjects were free to disengage from the task and leave the testing position without penalty. Engagement in the task was the sole criterion for inclusion. No additional exclusion criteria were applied, as none of the animals had been diagnosed with perceptual impairments that could have affected the visual or auditory processing of the stimuli. Of the eleven monkeys that initially approached the experimental setting, two individuals initiated the training phase but stopped participating before reaching the testing criterion and were therefore excluded from the study. All monkeys that proceeded to testing successfully completed the full set of test trials.

The protocol underwent review by the Mandai Wildlife Group Ethical Committee and was subsequently approved by the National University of Singapore Institutional Animal Care and Use Committee

### Training

Each monkey was trained individually using a shaping procedure. Importantly, no auditory stimuli were presented during training. In the initial shaping phase, monkeys were presented with a single metal cup placed on the table. The cup (8 cm in diameter) was covered with a squared lid (11 cm2) displaying a black shape composed of both round and spiky edges (**Fig. 2A**). The shape was identical to that used in a previous study on newly hatched chick (Loconsole et al., 2026), except that it was painted black, in line with Bouba–Kiki studies on humans (Bremner et al., 2013; Ćwiek et al., 2021a; Ramachandran & Hubbard, 2001). A small piece of banana was hidden underneath the lid. Monkeys learned to lift the lid to retrieve the food reward. Two different training shapes were generated based on the shapes used in the test phase; these were alternated across trials and randomly rotated by 180° to prevent reliance on low-level visual cues. Once subjects reliably lifted the lid to obtain the reward, they proceeded to a two-cup training phase. Two cups were placed simultaneously on the table: one cup was covered with a blank lid, and the other with a lid displaying the training shape. The food reward was always hidden beneath the lid displaying the training shape. The spatial position of the rewarded cup (left or right) was counterbalanced across trials and constrained so that it never appeared on the same side for more than two consecutive trials. A trial was considered correct when the monkey lifted the lid displaying the training shape and retrieved the food reward. Training continued until the subject reached a criterion of six consecutive correct trials, after which they advanced to the test phase.

**Fig. 2.**
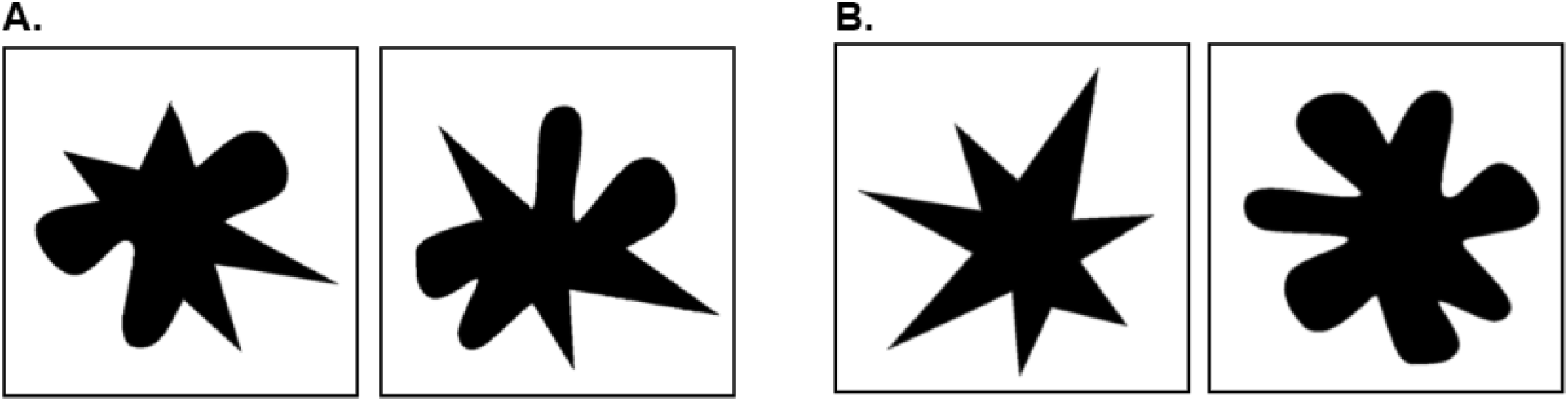
The training and testing stimuli. (A) Stimuli used at training. Each stimulus is obtained by blending the two testing shapes, so that training stimuli have both spiky and round edges. The stimuli were randomly alternated during testing trials. (B) Stimuli used at test. Both stimuli were presented simultaneously. These shapes were previously employed in studies on the Bouba-Kiki effect in humans and baby chicks.

### Test

Testing sessions were initiated in the same manner as training sessions, with the monkey voluntarily approaching the front of the home enclosure and positioning themselves near the experimenter and the table. To prevent interference from conspecifics, the tested individual was temporarily separated from other group members by closing a sliding door that restricted access to the testing area. In addition, a black curtain was positioned to block visual access to the testing setup, ensuring that other individuals could not observe the stimuli or the behavior of the tested subject. This procedure prevented both observational learning and incidental exposure to the round and spiky shapes prior to testing for the other monkeys.

Each test trial involved the simultaneous presentation of two metal cups, both covered with lids displaying a shape: one fully rounded and one fully spiky, identical to those used in previous studies with human participants and newly hatched chicks (**Fig. 2B**). Shapes were randomly rotated across trials to minimize the use of low-level visual cues. No food reward was hidden beneath either cup during test trials.

During each trial, either the sound “Bouba” or “Kiki” was played (at approximately 65dB) repeatedly from a speaker (Bose Soundlink mini II) positioned centrally underneath the table, equidistant from the two cups and concealed from the monkey’s view. The auditory stimuli were the same as those used in previous studies with humans (Ćwiek et al., 2021b) and chicks (Loconsole et al., 2026). There was no need to modify the auditory stimuli with respect to the original ones, as research on the genus Macaca shows that different species can successfully perceive and respond to auditory frequencies similar to humans (Jackson et al., 1999), as well as discriminate between different vowel sounds (Sinnott, 1989). The same applies to the visual stimuli: considering primates’ visual acuity (Iii et al., 2019; Srinivasan et al., 2015), we foresaw no impairment in shape discrimination. Notably, the visual stimuli used with chicks were painted orange for salience, as chickens are more responsive to this hue (Ham & Osorio, 2007), whereas we decided to maintain the original black coloration previously employed with humans and infants, as there is no evidence of a similar bias in macaques.

The spatial position of the two shapes (left/right) and the sound presented were pseudo-randomized across trials, with the constraint that the same condition never occurred on two consecutive trials. Throughout testing, the experimenter maintained a neutral posture and did not provide verbal, gestural, or positional cues. To avoid unintentionally guiding choices, the experimenter looked away from both the monkey and the stimuli during the decision phase. When necessary to engage attention and initiate interaction, the experimenter briefly tapped the two cups on the table and presented them clearly to the monkey, while carefully avoiding visual contact. A second experimenter remained at least two metres away from the apparatus and outside the monkey’s field of view. Once the first experimenter had shown the cups to the monkey and placed them on the table, the second experimenter started the audio playback, after which the monkey was allowed to make a choice. This procedure ensured that the first experimenter remained blind to the upcoming auditory stimulus while manipulating the visual stimuli.

A choice was defined as the lifting of one lid; only the first response in each trial was recorded, and the non-chosen cup was removed before the monkey could inspect it. A trial was classified as a non-response if the monkey left the testing position or failed to make a choice within 60 s. In such cases, the experimenter attempted to re-engage the monkey, and the same trial was repeated immediately afterwards. This happened for three monkeys (Joku, one trial; Sasha, one tiral; Silu, three trials; Shenaya, one trial. See Supplementary material **Data S1**). If two non-responses occurred consecutively, the testing session was terminated and resumed after a minimum interval of three hours, starting again from the last successfully completed trial. This happened for four monkeys (Shakira, session 1; Gaia, session 2; Shenaya, session 3; Kong, session 4). Each monkey completed a total of 24 test trials. To maintain motivation despite the absence of reward, test trials were organized into short blocks. After three consecutive unrewarded test trials, two rewarded “refresh” trials were administered, in which the training shape (both spiky and round edges) was presented against a blank lid, as in the training phase, followed by three additional unrewarded test trials. After six test trials, monkeys were given a break and were tested again after a minimum interval of one hour or, if necessary, on the following day. Before the start of each new test block (comprising three test trials, two refresh trials, and three test trials), monkeys completed a refresh session in which they were required to achieve six consecutive correct responses in the rewarded ambiguous-shape task. This criterion ensured that subjects remained engaged in the task and maintained adequate levels of attention and motivation before proceeding with further test trials.

All sessions were video-recorded, allowing for offline scoring of monkeys’ choices. Scoring was conducted independently by two experimenters from the video recordings. One of the coders was blind to the auditory condition of the trials to prevent observer bias.

### Quantification and statistical analysis

Data analyses were conducted using R software (R 4.4.0) (R Core Team, 2022). To test whether monkeys’ shape preferences were modulated by sound, we fitted a generalized linear mixed-effects model with a binomial error structure (R package: lme4 (Bates et al., 2015)) in which shape choice (round vs. spiky) was the response variable, background sound (Bouba vs. Kiki) was included as a fixed effect, and subject was included as a random intercept. The model exhibited a singular fit, possibly reflecting the small numbers of subjects (N = 9); however, residual diagnostics revealed no major violations of model assumptions (R package: DHARMa (Hartig & Lohse, 2022)). To assess whether the results depended on model specification, we compared the mixed-effects model to an alternative fixed-effects model in which subject was included as a factor. The two models did not differ significantly, χ^2^(7) = 7.16, p = .41, indicating that the random-intercept specification was adequate. To further assess robustness to the small sample size, we conducted leave-one-subject-out analyses in which the mixed-effects model was refit excluding each subject in turn. The effect of sound was highly stable across these analyses: odds ratios for Kiki relative to Bouba ranged from 0.14 to 0.21, and the direction of the effect did not change when any individual subject was removed. A Type II Wald χ^2^ test on the mixed-effects model showed that Sound significantly affected shape choice, χ^2^(1) = 35.14, p < .001. Estimated marginal means (R package emmeans(Lenth, 2022)) showed that in the Bouba trials monkeys were more likely than chance to choose the round shape (probability of round = 0.67, SE = 0.05, p < .001), whereas in the Kiki trials they preferentially chose the spiky shape (probability of round = 0.26, SE = 0.04, p < .001 (see Fig. 1D in main text). Consistent with these condition-specific effects, the odds of choosing the round shape were nearly six times higher in Bouba than in Kiki trials. (odds ratio = 5.95, 95% CI [∼3–12]).

Altogether, these results indicate that monkeys’ shape preferences are modulated by sound, consistent with cross-modal sound-symbolic mappings reported in other species.

## ACKNOWLEDGMENTS

Maria Loconsole’s work is funded by the European Union – NextGenerationEU and by the University of Padua under the 2023 STARS Grants@Unipd programme (project CROSS - Comparative Research Of Sound Symbolism). Elias Garcia-Pelegrin work’s is supported by a Mandai Wildlife Group Adjunct Principal Investigator position. We thank Mandai Wildlife Group for supporting our research and providing access to the macaque sample, with particular appreciation to Gail Laule.

## AUTHOR CONTRIBUTIONS

Conceptualization: M.L., C.X., E.G.P; Methodology: M.L., C.X., E.G.P; Investigation: M.L., C.X.; Resources: M.L., E.G.P.; Visualization: M.L., C.X.; Supervision: M.L., E.G.P.; Project administration, E.G.P.; Funding acquisition, M.L., E.G.P.; Writing—original draft: M.L.; Writing—review & editing: C.X., E.G.P.

## DATA, CODE, AND MATERIALS AVAILABILITY

The dataset generated during the study is available as Supplementary Material. Additional information supporting the findings of this study is available from the corresponding authors upon reasonable request.

## REFERENCES

Bates, D., Mächler, M., Bolker, B., & Walker, S. (2015). Fitting Linear Mixed-Effects Models Using lme4. Journal of Statistical Software, 67, 1–48. 10.18637/jss.v067.i01

Bremner, A. J., Caparos, S., Davidoff, J., de Fockert, J., Linnell, K. J., & Spence, C. (2013). “Bouba” and “Kiki” in Namibia? A remote culture make similar shape–sound matches, but different shape–taste matches to Westerners. Cognition, 126(2), 165–172. 10.1016/j.cognition.2012.09.007

Brunetti, R., Indraccolo, A., Del Gatto, C., Spence, C., & Santangelo, V. (2018). Are crossmodal correspondences relative or absolute? Sequential effects on speeded classification. Attention, Perception, & Psychophysics, 80(2), 527–534. 10.3758/s13414-017-1445-z

Ćwiek, A., Fuchs, S., Draxler, C., Asu, E. L., Dediu, D., Hiovain, K., Kawahara, S., Koutalidis, S., Krifka, M., Lippus, P., Lupyan, G., Oh, G. E., Paul, J., Petrone, C., Ridouane, R., Reiter, S., Schümchen, N., Szalontai, Á., Ünal-Logacev, Ö., … Winter, B. (2021a). The bouba/kiki effect is robust across cultures and writing systems. Philosophical Transactions of the Royal Society B: Biological Sciences, 377(1841), 20200390. 10.1098/rstb.2020.0390

Ćwiek, A., Fuchs, S., Draxler, C., Asu, E. L., Dediu, D., Hiovain, K., Kawahara, S., Koutalidis, S., Krifka, M., Lippus, P., Lupyan, G., Oh, G. E., Paul, J., Petrone, C., Ridouane, R., Reiter, S., Schümchen, N., Szalontai, Á., Ünal-Logacev, Ö., … Winter, B. (2021b). The bouba/kiki effect is robust across cultures and writing systems. Philosophical Transactions of the Royal Society B: Biological Sciences, 377(1841), 20200390. 10.1098/rstb.2020.0390

Emery, N. J., & Clayton, N. S. (2005). Evolution of the avian brain and intelligence. Current Biology, 15(23), R946–R950. 10.1016/j.cub.2005.11.029

Fort, M., & Schwartz, J.-L. (2022). Resolving the bouba-kiki effect enigma by rooting iconic sound symbolism in physical properties of round and spiky objects. Scientific Reports, 12(1), 19172. 10.1038/s41598-022-23623-w

Ghazanfar, A. A., Neuhoff, J. G., & Logothetis, N. K. (2002). Auditory looming perception in rhesus monkeys. Proceedings of the National Academy of Sciences, 99(24), 15755–15757. (world). 10.1073/pnas.242469699

Ham, A. D., & Osorio, D. (2007). Colour preferences and colour vision in poultry chicks. Proceedings. Biological Sciences, 274(1621), 1941–1948. 10.1098/rspb.2007.0538

Hartig, F., & Lohse, L. (2022). DHARMa: Residual Diagnostics for Hierarchical (Multi-Level / Mixed) Regression Models (Version 0.4.6) [Computer software]. https://cran.r-project.org/web/packages/DHARMa/index.html

Iii, W. H. R., Zhang, K. M., Karsolia, A., Engles, M., & Burke, J. (2019). Comparison of contrast sensitivity in macaque monkeys and humans. Visual Neuroscience, 36, E008. 10.1017/S0952523819000051

Jackson, L. L., Heffner, R. S., & Heffner, H. E. (1999). Free-field audiogram of the Japanese macaque (Macaca fuscata). The Journal of the Acoustical Society of America, 106(5), 3017–3023. 10.1121/1.428121

Jarvis, E. D., Güntürkün, O., Bruce, L., Csillag, A., Karten, H., Kuenzel, W., Medina, L., Paxinos, G., Perkel, D. J., Shimizu, T., Striedter, G., Wild, J. M., Ball, G. F., Dugas-Ford, J., Durand, S. E., Hough, G. E., Husband, S., Kubikova, L., Lee, D. W., … Butler, A. B. (2005). Avian brains and a new understanding of vertebrate brain evolution. Nature Reviews Neuroscience, 6(2), 151–159. 10.1038/nrn1606

Köhler, W. (1970). Gestalt Psychology: An Introduction to New Concepts in Modern Psychology.Liveright.

Lenth, R. (2022). emmeans: Estimated Marginal Means, aka Least-Squares Means. https://CRAN.R-project.org/package=emmeans

Lockwood, G., & Dingemanse, M. (2015). Iconicity in the lab: A review of behavioral, developmental, and neuroimaging research into sound-symbolism. Frontiers in Psychology,6. https://www.frontiersin.org/articles/10.3389/fpsyg.2015.01246

Loconsole, M., Benavides-Varela, S., & Regolin, L. (2026). Matching sounds to shapes: Evidence of the bouba-kiki effect in naïve baby chicks. Science, 391(6787), 836–839. 10.1126/science.adq7188

Loconsole, M., Pasculli, M. S., & Regolin, L. (2021). Space-luminance crossmodal correspondences in domestic chicks. Vision Research, 188, 26–31. 10.1016/j.visres.2021.07.001

Loconsole, M., & Regolin, L. (2023). Here I am, why don’t you answer me? Sensitivity to social responsiveness in domestic chicks. iScience, 26(1), 105863. 10.1016/j.isci.2022.105863

Loconsole, M., Stancher, G., & Versace, E. (2023). Crossmodal association between visual and acoustic cues in a tortoise (Testudo hermanni). Biology Letters, 19(7), 20230265. 10.1098/rsbl.2023.0265

Ludwig, V. U., Adachi, I., & Matsuzawa, T. (2011). Visuoauditory mappings between high luminance and high pitch are shared by chimpanzees (Pan troglodytes) and humans. Proceedings of the National Academy of Sciences, 108(51), 20661–20665. 10.1073/pnas.1112605108

Margiotoudi, K., Allritz, M., Bohn, M., & Pulvermüller, F. (2019). Sound symbolic congruency detection in humans but not in great apes. Scientific Reports, 9(1), 12705. 10.1038/s41598-019-49101-4

Margiotoudi, K., Bohn, M., Schwob, N., Taglialatela, J., Pulvermüller, F., Epping, A., Schweller, K., & Allritz, M. (2022). Bo-NO-bouba-kiki: Picture-word mapping but no spontaneous sound symbolic speech-shape mapping in a language trained bonobo. Proceedings of the Royal Society B: Biological Sciences, 289(1968), 20211717. 10.1098/rspb.2021.1717

Mondloch, C. J., & Maurer, D. (2004). Do small white balls squeak? Pitch-object correspondences in young children. Cognitive, Affective, & Behavioral Neuroscience, 4(2), 133–136. 10.3758/CABN.4.2.133

Ozturk, O., Krehm, M., & Vouloumanos, A. (2013). Sound symbolism in infancy: Evidence for sound-shape cross-modal correspondences in 4-month-olds. Journal of Experimental Child Psychology, 114(2), 173–186. 10.1016/j.jecp.2012.05.004

R Core Team. (2022). R: A Language and Environment for Statistical Computing. https://www.R-project.org/

Ramachandran, V. S., & Hubbard, E. M. (2001). Synaesthesia—A window into perception, thought and language. Journal of Consciousness Studies, 8(12), 3–34.

Seyfarth, R. M., Cheney, D. L., & Marler, P. (1980). Vervet monkey alarm calls: Semantic communication in a free-ranging primate. Animal Behaviour, 28(4), 1070–1094. 10.1016/S0003-3472(80)80097-2

Sinnott, J. M. (1989). Detection and discrimination of synthetic English vowels by Old World monkeys (Cercopithecus, Macaca) and humans. The Journal of the Acoustical Society of America, 86(2), 557–565. 10.1121/1.398235

Spence, C. (2011). Crossmodal correspondences: A tutorial review. Attention, Perception, & Psychophysics, 73(4), 971–995. 10.3758/s13414-010-0073-7

Spence, C. (2019). On the Relative Nature of (Pitch-Based) Crossmodal Correspondences.Multisensory Research, 32(3), 235–265. 10.1163/22134808-20191407

Spence, C. (2022). Exploring Group Differences in the Crossmodal Correspondences.Multisensory Research, 35(6), 495–536. 10.1163/22134808-bja10079

Spence, C., & Deroy, O. (2012). Crossmodal correspondences: Innate or learned? I-Perception,3(5), 316–318. 10.1068/i0526ic

Srinivasan, S., Carlo, C. N., & Stevens, C. F. (2015). Predicting visual acuity from the structure of visual cortex. Proceedings of the National Academy of Sciences, 112(25), 7815–7820. 10.1073/pnas.1509282112

